# miRNA-mediated regulation of lineage plasticity in YY1 knockout pro-B cells

**DOI:** 10.64898/2026.09.15.751727

**Authors:** Nasreen Bano, Sarah Naiyer, Sulagna Sanyal, Suchita Hodewadekar, Michael L. Atchison

## Abstract

Yin Yang 1 (YY1) is a multifunctional transcription factor involved in chromatin organization and gene regulation. While the mechanisms by which YY1 controls B cell lineage development have been explored at the transcriptional level, little is known at the post-transcriptional level. To address this, we performed integrated analyses combining bulk transcriptomics, miRNA profiling, and single-cell transcriptomics to identify YY1-dependent regulatory networks at the pro-B cell stage. YY1 knockout (KO) in pro-B cells leads to reduced B-lineage commitment and activation of genes associated with alternative hematopoietic fates. Our integrative analyses suggest that these changes are organized in part through coordinated miRNA-mRNA regulatory networks. In wild-type (WT) pro-B cells, miRNAs such as miR-10a-5p, miR-674-5p, and miR-21a-5p maintain B cell identity by suppressing RNAs linked to other lineages. In contrast, in YY1 KO pro-B cells increased expression of miRNAs including miR-15b-5p, miR-342-5p, and miR-34a-5p target pathways required for B cell development, proliferation, and survival thus reducing B lineage identity. Together, these findings suggest that YY1 preserves B cell identity through coordinated regulation of transcriptional and post-transcriptional programs, with YY1-dependent miRNA networks contributing to the maintenance of B cell-associated gene expression and restriction of alternative lineage programs. Among these, miR-15b-5p emerged as a central regulator predicted to regulate Bcl6b and E2f family transcription factors that control proliferative and transcriptional programs critical for early B cell development. At single-cell resolution, YY1 KO pro-B cells grown on OP9-DL4 feeders that provide Notch signaling displayed marked transcriptional heterogeneity and loss of pro-B lineage commitment. Pseudotime analysis revealed a branched developmental landscape, with cells diverging toward monocyte, macrophage, and dendritic cell fates rather than a consistent B cell differentiation trajectory. Collectively, these findings establish YY1 as a central regulator of lineage commitment integrating transcriptional and miRNA-mediated post-transcriptional mechanisms to regulate B cell identity.

## Introduction

Yin Yang 1 (YY1) is a multifunctional transcription factor with essential roles in embryogenesis, cellular proliferation, DNA replication, gene expression, and lineage-specific differentiation across multiple biological systems [1, 2]. Functionally, YY1 acts as both a transcriptional activator and repressor by recruiting to DNA diverse regulatory complexes, including co-activators, co-repressors, and Polycomb group (PcG) proteins [3–6]. In addition, YY1 directly interacts with core components of the basal transcription machinery, such as TATA-binding protein (TBP), TBP-associated factors (TAFs), and transcription factor IIB, further underscoring its central role in gene regulation [7]. Beyond direct transcriptional control, YY1 has emerged as a key regulator of higher-order chromatin organization, contributing to enhancer-promoter communications and three-dimensional genome architecture [8–12].

Hematopoiesis provides a well-defined system for studying lineage commitment. Hematopoietic stem cells differentiate into lymphoid (B and T cells), myeloid (macrophages, dendritic cells, etc.), and erythroid progenitors. Each hematopoietic lineage is controlled by multiple transcription factors and chromatin remodeling proteins, which also impact three-dimensional chromatin architecture [13]. B lymphocytes are crucial components of the adaptive immune response and are responsible for humoral immunity in mammals [14]. The early stages of B cell development are regulated by at least 10 distinct transcription factors, with E2A, *Ebf1*, and *Pax5* being especially crucial for B-lineage commitment and differentiation [15]. Previous studies have established that YY1 is also crucial for early B cell development, with conditional deletion at the pro-B stage leading to developmental arrest and profound disruption of B-lineage programs [2, 16]. Our recent work showed that YY1 knockout (KO) in pro-B cells leads to unusual lineage plasticity, enabling these cells to develop into alternative hematopoietic lineages [2]. These findings position YY1 not only as a transcriptional activator of B cell identity genes but also as a critical gatekeeper that restricts alternative lineage potential. Loss of YY1 may therefore create a permissive developmental state in which pro-B cells become more responsive to alternative lineage-promoting cues.

While transcriptional regulation is central to lineage commitment, post-transcriptional mechanisms, particularly those mediated by microRNAs (miRNAs), a class of small non-coding RNAs, provide an additional layer of control to modify gene expression networks. These small non-coding RNAs regulate gene expression by promoting mRNA degradation or inhibiting translation and thereby shaping diverse biological processes including differentiation, proliferation, and apoptosis [17–21]. Dysregulation of miRNA expression has been implicated in developmental abnormalities and immune disorders [22]. However, the extent to which YY1 coordinates miRNA-mediated regulatory networks to enforce B cell lineage fidelity remains poorly understood. To address this gap, we performed an integrated multi-omics analysis combining bulk RNA-seq, miRNA-seq, and single-cell RNA-seq (scRNA-seq) to define YY1-dependent transcriptional and post-transcriptional regulatory programs in pro-B cells. Specifically, we found that miRNAs enriched in wild-type pro-B cells, including miR-181b-5p, miR-17-5p, miR-21a-5p, miR-142a-5p target transcripts associated with alternative lineage pathways such as Notch signaling, T-cell receptor signaling, and stem cell pluripotency, thereby reinforcing B-lineage commitment. In addition, we identified a subset of miRNAs (miR-15b-5p, miR-342-5p, miR34a-5p, miR-103-5p, miR-150-5p, miR-151-rp, miR-423-5p, miR103-2-5p, and miR103-1-5p) that are induced upon YY1 KO in pro-B cells. We found miR-15b-5p, in particular, down regulates transcripts important for B lineage development. These findings suggest that YY1-dependent regulation of miR-15b-5p may contribute to the destabilization of B cell identity following YY1 loss. Integration with single-cell transcriptomic data further reveal that disruption of YY1-centered regulatory networks results in marked transcriptional changes enabling YY1 KO pro-B cells to acquire multilineage potential.

Collectively, our findings establish that YY1-dependent maintenance of B cell identity involves both transcriptional regulation and miRNA-mediated post-transcriptional control. These results reveal an additional regulatory layer through which YY1 contributes to lineage fidelity and developmental stability.

## Methods

### Data retrieval and processing of bulk RNA-sequencing datasets

The raw RNA-seq fastq triplicate files of murine WT pro-B cells (B) and YY1-null pro-B cells (Y) were retrieved from the National Center for Biotechnology Information-Sequence Read Archive (NCBI-SRA) with accession number PRJNA297235. The quality of fastq raw reads was checked using FastQC v.0.12.0 [23]. Low-quality reads were eliminated using Trimmomatic 0.36 [24] at the default parameter. Unsupervised clustering was performed among the replicate samples using the factoextra package in RStudio [25]. Hierarchical clustered dendrogram analysis was also performed using the complete Pearson method in R.

### Identification of differentially expressed genes in transcriptome data

Filtered reads were mapped against the GRCm38 (mm10) mouse genome using STAR [26]. The mm10 database was retrieved from the Ensembl database (http://useast.ensembl.org/Mus_musculus/Info/Index). To quantify transcript abundance of mRNAs, StringTie v2.1.2 [27] was used. Read counts were then subjected to edgeR/Bioconductor R package v.4.0.2 [28] to normalize read counts and identify differentially expressed mRNAs among the pro-B cell (B) and YY1-null pro-B cell (Y) samples. Differential expression of pro-B cell (B) and YY1-null pro-B cell (Y) transcripts was determined based on the criteria of | log2(Fold Change) | ≥ 1 with p-value ≤ 0.05 for upregulated genes and |log2(Fold Change) | ≤ 1 with p-adjusted value ≤ 0.05 for downregulated genes. Gene annotation of differentially expressed genes (DEGs) was carried out with the GRCm38 (mm10) mouse genome. 3D PCA analyses were performed using the factoextra package in R. Volcano plots of differentially expressed mRNAs (DEmRNAs) were generated using ggplot2 [29].

### Hierarchical clustering of DEGs

Hierarchical clustering of log_2_-transformed expression data was carried out using the fuzzy c-means algorithm in RStudio using the Mfuzz package [30]. In contrast to rigid clustering methods like K-means, this algorithm somewhat mitigates the impact of noise on clustering outcomes, and it effectively defines the genes and relationships between clusters. All identified differentially expressed genes were divided into six different clusters according to their expression pattern in pro-B cell (B) and YY1-null pro-B (Y) cell samples.

### scRNA-seq

scRNA-seq data from *yy1f/f* and *yy1^f/f^ Mb1-CRE* mice grown on OP9-DL4 feeder cells is described in Banerjee 2024 [2].

### RNA Extraction for miRNA-seq

Total RNA was extracted from the mouse bone marrow cells. Fluorescence-activated cell-sorted (FACS) cells were collected directly in TRIzol using the Direct-zol RNA Purification Kit according to the manufacturer’s instructions. This method allows efficient purification of total RNA, including small RNAs, directly from TRIzol lysates without phase separation. RNA concentration was quantified using a Qubit 4 Fluorometer (Thermo Fisher Scientific) with the Qubit RNA HS Assay Kit. RNA integrity was assessed using an Agilent 4200 TapeStation system. Only samples with RNA Integrity Number (RIN) ≥ 7.5 and clear small RNA peaks were used for library preparation.

### Small RNA library preparation and sequencing

Small RNA libraries were prepared using the QIAseq miRNA Library Kit (Qiagen) following the manufacturer’s protocol. Briefly, 3′ and 5′ adapters were ligated to the small RNA molecules, followed by reverse transcription and PCR amplification (15 cycles) using indexed primers to allow multiplex sequencing. Size selection was performed to enrich for miRNA-derived inserts. Final library sizes ranged between 190-211 bp. Library concentration and quality were evaluated using the Agilent TapeStation system. Libraries were pooled in equimolar amounts and sequenced on an Illumina NextSeq 2000 platform using single-read mode with 75 bp read length (SR75). Each sample generated approximately 10-15 million reads. The raw reads generated in the current study were submitted to NCBI-SRA with BioProject PRJNA1510790.

### Bioinformatics quality control and pre-processing of miRNA-seq

miRNA-seq data were obtained from sorted pro-B cells from YY1 WT (*yy1f/f*) and YY1 KO (*yy1f/f Mb1Cre*) mice. Raw sequencing reads were assessed for quality using FastQC (v0.12.1) to evaluate per-base sequence quality, GC content, sequence duplication levels, and overrepresented sequences. Adapter sequences and low-quality bases were trimmed using Cutadapt (v4.4) with the following parameters: minimum read length of 18 bp, Phred quality score threshold of 30, and trimming of Illumina 3′ adapters. Reads containing ambiguous bases (N) were discarded. After trimming, read quality was re-evaluated using FastQC to confirm improvement. The length distribution of processed reads was assessed to confirm enrichment of 18-26 nucleotide sequences characteristic of mature miRNAs. Further quality control included estimation of sequencing depth and saturation analysis. Only samples with sufficient read depth and high-quality metrics (average Phred score > 28 post-trimming and > 70% reads retained after filtering) were retained for downstream analysis. Across all samples, miRNA sequencing yielded 12.3-13.8 million raw reads per replicate. After quality filtering and adaptor trimming, 88-91% of unmapped reads were used to identify mature miRNAs (Table S1).

### miRNA identification and quantification

The miRDeep2 v.0.1.3 program identified and quantified miRNAs using default parameters [31]. Filtered RNA sequences were used against the reference mouse genome GRCm38 (mm10) for the identification of miRNAs. The GRCm38 (mm10) genome was retrieved from the Ensembl database and compared with the miRBase database version 22.1 [32]. Filtered sequences were aligned to the mouse reference genome GRCm38 (mm10), and mature miRNAs (18-25 nt) were identified using miRDeep2 based on alignment to known miRNA sequences and a significant randfold p-value. miRDeep2 was also used for the quantification of miRNA expression levels in different samples. The miRNA gene expression matrix data were then subjected to DESeq2 [33] to identify the differentially expressed miRNAs among the WT proB (*yy1f/f*) and YY1KO pro-B (*yy1f/f Mb1CRE*) cell samples. Significantly differentially expressed miRNAs were determined based on the criteria of |log_2_(Fold Change) | ≥ 2 and FDR (adjusted P value ≤ 0.05).

### Target identification of miRNAs and their annotations

To understand the functions of differentially expressed miRNAs, their putative targets were predicted using miRDB (https://mirdb.org/) [34]. A target prediction score ≥ 60 was used to identify high-confidence putative targets. In miRDB, the prediction score ranges from 50 to 100, with higher scores indicating greater confidence that the predicted miRNA-target interaction is real and functionally relevant. Functional annotations of the predicted targets were performed using the mouse genome in miRDB. To further refine the selection of target transcripts for downstream enrichment analysis, a more stringent prediction-score threshold of ≥ 80 was applied.

### Enrichment analysis

To study the putative functions of the identified differentially expressed genes, gene ontology (GO) analysis was performed classifying genes into cellular component (CC), molecular function (MF), and biological process (BP) categories. GO enrichment analysis was performed using org.Mm.eg.db keytype and the enrichplot package in RStudio with < 0.05 adjusted p-value cutoff for significantly enriched terms [35]. GO visualization was performed using the ggplot2 package in R. To understand the enriched pathways for significant DEmRNAs and DEmiRNAs, Kyoto encyclopedia of genes and genomes (KEGG) analysis was performed. KEGG pathways analysis was employed using the clusterProfiler package with the number of permutations (nPerm) set to 10,000, org.Mm.eg.db and pAjustMethod set to “BH”. P-value ≤ 0.05 was set for significant KEGG pathways [36].

### scRNA-seq data analysis

The scRNA-seq raw data were retrieved from the National Center for Biotechnology Information Sequence Read Archive (NCBI-SRA) with bioproject accession number PRJNA1073566 [12], which comprises four samples: (A) YY1 WT *yy1f/f* undifferentiated, (B) YY1 KO *yy1f/f Mb1-CRE* undifferentiated (A and B cells directly isolated from bone marrow), (C) YY1 WT *yy1f/f* differentiated by culture on OP9-DL4 feeder cells for 14 days, and (D) YY1 KO *yy1f/f Mb1-CRE* cells differentiated by culture on OP9-DL4 feeders for 14 days [2]. The conversion of YY1-null pro-B cells into T-like cells on OP9-DL4 feeder layers requires approximately three weeks, resulting in a population comprising over 90% CD25⁺ Thy1⁺ cells [2]. However, harvesting cells at an intermediate stage (14 days) captures key transitional states including monocytes, dendritic cells, NK cells, and stem cells [2]. Thus, pro-B cells were isolated from *yy1^f/f^* and *yy1^f/f^ mb1-CRE* mice and cultured on OP9-DL4 cells with IL7, Flt3L, and SCF until 3%-5% of the *yy1^f/f^* mb1-CRE pro-B population had become CD25+ Thy1+ (as expected, cells from wild-type *yy1^f/f^* mice remained negative). Cells from both genotypes cultured on OP9-DL4 feeders were harvested and subjected to single-cell RNA sequencing (scRNA-seq), along with pro-B cells directly isolated from *yy1^f/f^* and *yy1^f/f^ Mb1CRE* mice [2].

The raw gene expression matrices were generated by the cell ranger software (10X Genomics, v8.0.0). The mkfastq command was used for demultiplexing to generate FASTQ files. After demultiplexing, the resulting fastq files were aligned against the mm10 reference genome and feature-barcode matrices were generated with cellranger count [37]. For each sample, the recovered-cell parameter was set to 10,000 cells, which we expect to recover for each library. The output-filtered gene count matrices were analyzed by R (v4.4.1) using the Seurat package (v5.3.0) [38]. Different thresholds were chosen because of their distinct distribution in each sample to get good-quality cells [39]. The unique molecular identifier (UMI) count tables were imported with the Read10X_h5 function, and Seurat objects were created for each sample. We filtered out cells that had more than 5% of mitochondrial genes (Figure S4). Samples with similar conditions were merged, data were normalized with SCTransform, which also regressed out the effect of library size, and most variable genes were detected by the FindVariableFeatures function. All samples were integrated using Seurat functions FindIntegrationAnchors and IntegrateData. After integration, the data were scaled using ScaleData, regressing out the effects ofUMI counts and mitochondrial gene content. Principal component analysis (PCA) was then performed using RunPCA for dimensionality reduction (30 PCs). For clustering, the FindNeighbors (20 PCA) and FindClusters (resolution 0.6) functions were used. FindAllMarkers was used to compare a cluster against all other clusters to identify the marker genes. For each cluster, the minimum required average log fold change in gene expression was set to 0.25, and the minimum percent of cells that must express genes in either cluster was set to 25%. Cell types were annotated with ImmGen reference data using the SingleR (v2.10.0) package. Marker gene expression was visualized with FeaturePlot in RStudio. Pseudotime trajectory analysis was performed using Monocle3 (v1.4.27). UMAP was used for dimensionality reduction, and a principal trajectory graph was inferred in UMAP space using the Monocle3 learn_graph() function with use_partition = TRUE. A principal graph node within the population classified by SingleR as stem/progenitor-like was manually designated as the root using order_cells(). The root was selected based on the progenitor-associated transcriptional identity of the corresponding cells, providing a biologically informed reference point for pseudotime ordering.

### Statistical analysis

All the relative expression graphs were plotted by the use of GraphPad Prism for Mac OS version 10.2.0 (GraphPad Software, San Diego, CA, USA). Two-way analysis of variance (ANOVA) was used for statistical analysis. The value of P<0.05 was counted as statistically significant.

## Results

### YY1 KO alters gene expression patterns in pro-B cells

We analyzed RNA-seq data from WT pro-B cells (B) and YY1 KO pro-B cells (Y), each with three biological replicates (B1, B2, B3, Y1, Y2, Y3) [40]. Low-quality reads were filtered and remaining reads were mapped to the mm10 mouse genome. The resulting mapped read counts were then normalized and utilized to identify differentially expressed genes (DEGs) in WT pro-B cells (B) compared to YY1 KO pro-B cells (Y). Quality-control analyses using Pearson correlation, hierarchical clustering, and principal component analysis revealed clear overall separation between YY1 WT and YY1 KO samples (Fig. S1A-C). B1 and B2 were strongly correlated (r = 0.94), whereas B3 showed lower correlations with B1 and B2 (r = 0.54 and 0.52, respectively). Nevertheless, B3 clustered with the other YY1 WT samples in the hierarchical clustering and PCA analyses, supporting its inclusion in the downstream analysis.

A total of 5,031 unique genes were expressed in YY1 WT pro-B cells (26.82%) and 70 unique genes were uniquely expressed in YY1 KO pro-B cells (0.37%), with 13,657 genes (72.80%) being common to both groups (Fig. 1A). Heatmap representation of the top 15 B lineage or the top 15 T cell and other hematopoietic lineage genes expressed in YY1 WT pro-B cells (B) and YY1 KO pro-B cells (Y) is shown in Figure 1B. These data show that YY1 knockout reduces expression of numerous B lineage genes but increases the expression of alternative lineage genes (red arrows indicate key B or alternative hematopoietic lineage genes). This is consistent with our recent work showing YY1 KO pro-B cells are capable of adopting other alternative hematopoietic fates if placed in specific conditions that stimulate the generation of alternative lineages [2]. We identified a total of 1,233 significantly differentially expressed genes comparing YY1 KO pro-B (Y) vs YY1 WT pro-B (B) samples. Among these, 601 genes were up-regulated (log2FC ≥ 1 and p-adj ≤ 0.05), while 632 genes were down-regulated (log2FC ≤ −1 and p-adj ≤ 0.05). Volcano plot analysis (Fig. 1C) highlights six highly expressed genes associated with B cells that are significantly down-regulated upon YY1 KO in pro-B cells (*Blk, VpreB1, VpreB2, Igl1, Blnk, Cd79b*), and six highly expressed genes associated with alternative hematopoietic lineages that are significantly increased in expression upon YY1 KO (*Bcl6, Notch2, Cish, Cd300lf, Clec4d, Sema4a*). Thus, YY1 has a significant impact on expression of B lineage genes, as well as repression of alternative lineage genes.

**Figure 1.**
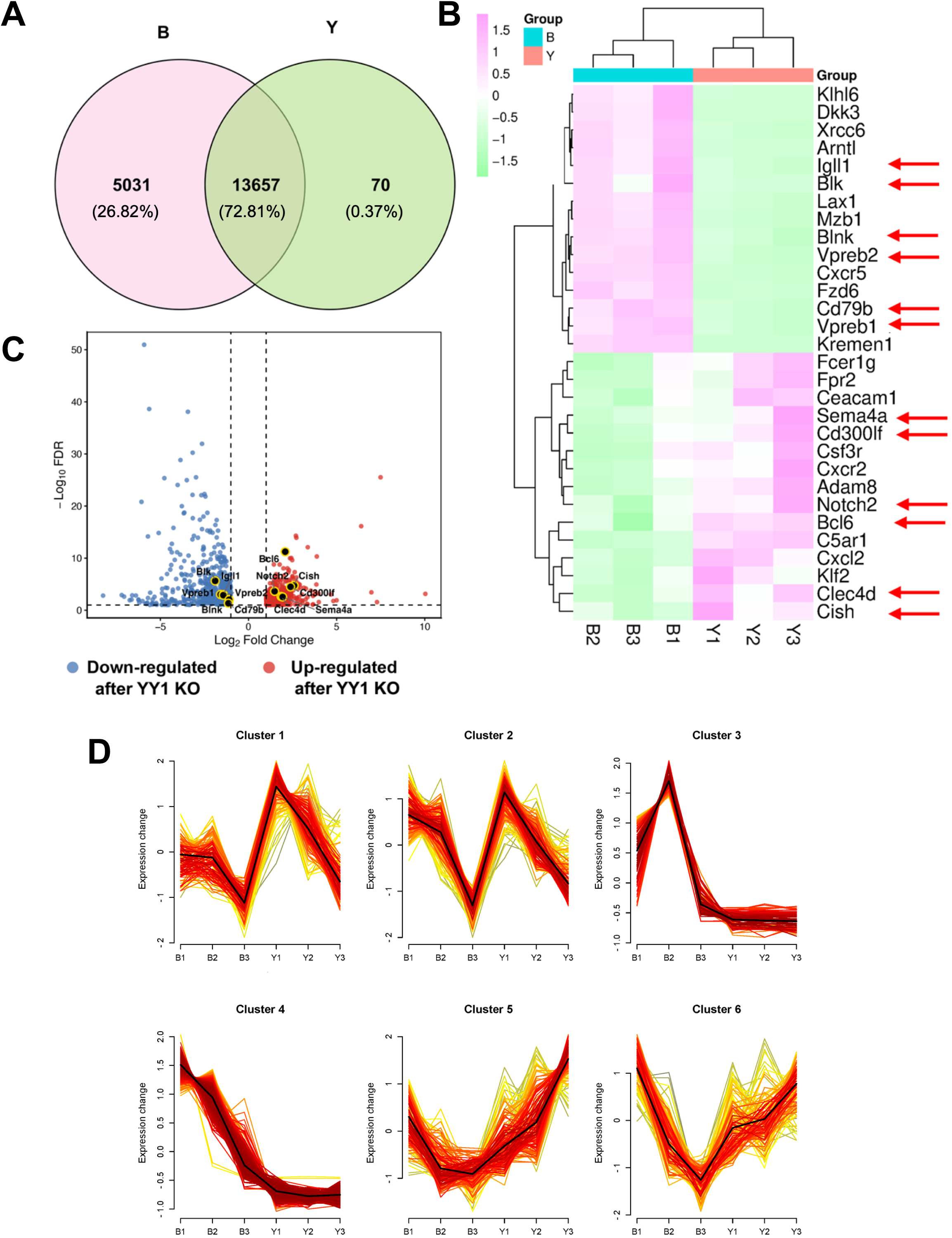
Transcriptomic differences between wild-type (WT) and YY1 KO pro-B cells. **(A)** Venn diagram showing the overlap of expressed transcripts between WT pro-B cells (B) and YY1 KO pro-B cells (Y). Numbers indicate transcript counts unique to each population or shared between them. **(B)** Hierarchical clustering heatmap of the top 15 B cell, or top 15 T cell and other hematopoietic lineage differentially expressed mRNAs in either YY1 WT or KO pro-B cells. Color scale (green to pink) indicates relative expression levels from low to high, respectively. Red arrows indicate key transcripts that are differentially expressed in either WT or YY1 KO pro-B cells **(C)** Volcano plot of RNA-seq data illustrating differentially expressed mRNAs (DEmRNAs) between WT (B) and YY1 KO (Y) pro-B cells. Red dots represent significantly up-regulated genes, and blue dots represent significantly down-regulated genes in YY1 KO pro-B cells (|log₂ fold change| ≥ 1 and adjusted P ≤ 0.05). The top six most significantly differentially expressed genes are labeled and correspond to the transcripts indicated by red arrows in Panel B. **(D)** Clustering of differentially expressed genes based on expression dynamics between wild-type and YY1 KO pro-B cells. Differentially expressed mRNAs identified between wild-type (WT) pro-B cells (B1–B3) and YY1 KO pro-B cells (Y1–Y3) were subjected to fuzzy c-means clustering using the Mfuzz package according to their expression patterns across the six samples (three biological replicates per genotype). Six distinct clusters were identified, each depicting a unique dynamic expression profile. The red and yellow lines represent the expression trajectories of individual genes, while the bold black lines indicate the average (centroid) expression pattern for all genes within each cluster. Cluster sizes are indicated in parentheses: Cluster 1 (135 genes), Cluster 2 (185 genes), Cluster 3 (90 genes), Cluster 4 (542 genes), Cluster 5 (164 genes), and Cluster 6 (117 genes). This analysis highlights coordinated transcriptional programs that are differentially regulated in YY1 KO pro-B cells.

The biological impact of a particular gene can depend upon both the cell-specificity of expression as well as the overall level of gene expression within a cell. A two-fold difference in expression of a rare transcript may yield little biological impact, whereas a 2-fold difference in a more abundant transcript may yield strong consequences. Therefore, we used the fuzzy c-means algorithm from the Mfuzz package [30], taking into account the level of gene expression in defining comparative groupings between samples. In contrast to rigid clustering methods such as K-means, this algorithm somewhat mitigates the impact of noise on clustering outcomes and effectively defines genes and relationships between clusters [41]. Using this approach, expression patterns of all differentially expressed genes in WT (B) and YY1 KO (Y) pro-B cells were represented in 6 clusters, each with distinct differentially expressed genes (Figure 1D). Genes with higher expression in wild-type pro-B cells (B) compared to YY1 KO pro-B cells (Y) samples were grouped into clusters 3 and 4. On the other hand, genes that were more highly expressed in YY1KO pro-B cells (Y) compared to WT pro-B cells (B) grouped into cluster 5 (Figure 1D). Clusters 1,2, and 6 did not yield a clear perspective on gene expression patterns in the triplicate samples. We therefore excluded these samples from further analyses and focused on evaluation of the genes in clusters 3, 4, and 5.

### YY1 KO drives distinct expression patterns across clusters

Our results show that clusters 3 and 4 contain genes with higher expression in WT pro-B cells compared to YY1 KO pro-B cells, whereas cluster 5 contains genes with higher expression in YY1 KO pro-B cells (Fig. 1D). Gene ontology and KEGG analyses (BP, Biological process; CC, Cellular component; MF, Molecular function) revealed distinct results for each cluster. Cluster 3 showed enrichment of diverse biological processes and pathways, including cell-cell adhesion, action potential, MAPK signaling, and cAMP signaling, but these pathways were associated with low gene counts and less significant P-values. Therefore, Cluster 3 was excluded from further analyses.

Cluster 4 genes were expressed at higher levels in WT pro-B cells and were strongly enriched for B lineage-associated processes and B cell pathway enrichments containing substantial numbers of genes (ranging from 10 to 25) and significant statistical p-values (Table S2, cluster 4 tab). Cluster 4 genes also revealed both B cell receptor signaling and Th1 and Th2 cellular differentiation pathways, in addition to immunodeficiency, and various metabolic, amino acid, and oxidative phosphorylation pathways (Fig. 2A and Table S2, cluster 4 tab). Representative genes included key B cell receptor signaling components (*Blk, Nfkbie, Cd72, Rasgrp3, Cd79b, and Blnk*) and Th1 and Th2 differentiation genes (*H2-Ob, Nfkbie, Stat4, H2-Oa, Il12a, Prkcq, and Zap70*) (Fig. 2B). Expression of these genes was markedly reduced in YY1 KO pro-B cells (Fig. 2C and D). This is consistent with reduced B cell identity and increased lineage plasticity we previously observed upon YY1 deletion in pro-B cells [2].

**Figure 2.**
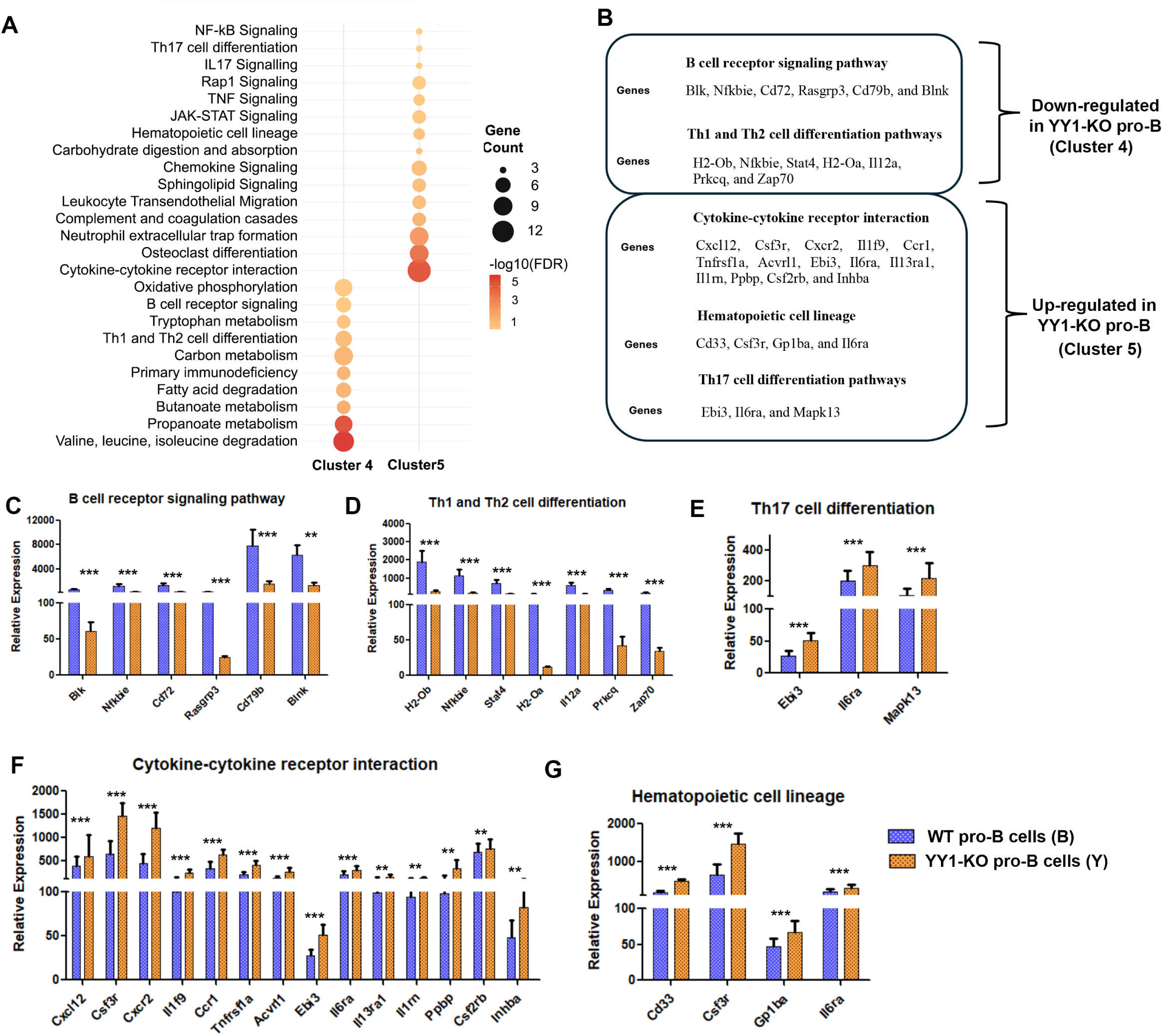
KEGG pathway enrichment analysis and validation of differentially expressed genes in WT and YY1 KO pro-B cells. **(A)** Bubble plot showing KEGG pathway enrichment of differentially expressed genes in clusters 4 and 5. Circle size represents the number of enriched genes, and circle color indicates the false discovery rate (FDR)-values. **(B)** Representative genes associated with selected enriched pathways. Cluster 4 genes are downregulated and cluster 5 genes are upregulated in YY1 KO pro-B cells relative to YY1 WT pro-B cells. **(C-D)** Relative mRNA expression of cluster 4 genes involved in the B cell receptor signaling pathway (C) or Th1 and Th2 cell differentiation (D) **(E-G)** mRNA expression of cluster 5 genes involved in Th17 cell differentiation (E), cytokine-cytokine receptor interaction (F), or hematopoietic cell lineage (G) pathways. Data are from three independent biological replicates. The error bars represent standard deviation of the mean. The asterisks indicate significant differences: *P < 0.01, **P < 0.001, ***P < 0.0001.

Cluster 5 genes were upregulated in YY1 KO pro-B cells and showed enrichment for alternative hematopoietic lineage genes and pathways. These pathways consisted of much smaller numbers of genes involved in B lineage processes, but increased numbers of genes involved in alternative hematopoietic lineages (Table S2, cluster 5 tab). Alternative lineage pathways included cytokine-cytokine receptor interaction, hematopoietic cell lineage, JAK-STAT signaling, IL-17, TNF, Th17, NF-κB, and Rap1 pathways, as well as processes associated with myeloid leukocyte differentiation and activation (Figure 2A, Table S2, cluster 5 tab). Notably, this cluster also included pathways linked to osteoclast differentiation and neutrophil extracellular trap formation (Figure 2A). Key upregulated genes included those with cytokine and receptor interaction functions (*Cxcl12, Csf3r, Cxcr2, Il1f9, Ccr1, Tnfrsf1a, Acvrl1, Ebi3, Il6ra, Il13ra1, Il1rn, Ppbp, Csf2rb,* and *Inhba*), hematopoietic lineage markers (*Cd33, Csf3r, Gp1ba, and Il6ra*), Th17 differentiation (*Ebi3, Il6ra, and Mapk13*), and regulators of myeloid and alternative lineage functions (Fig. 2B, E, F, G). These findings indicate that YY1 loss leads to increased expression of alternative hematopoietic lineage programs, supporting the role of YY1 in restricting non-B cell fate potential in pro-B cells [2].

### YY1 KO alters miRNA gene expression

Micro-RNAs (miRNAs) can also influence lineage development by post-transcriptionally reducing specific mRNA populations [42]. Therefore, we sought to determine if YY1 KO also had an impact on expression of various miRNAs. High-quality miRNA-seq data were obtained from sorted pro-B cells from YY1 WT (*yy1f/f*) and YY1 KO (*yy1f/f Mb1Cre*) mice. To further evaluate the quality and reproducibility of the miRNA-seq dataset, we performed correlation analyses across biological replicates. Scatter plots of log₂-transformed normalized miRNA expression values demonstrated high reproducibility between replicates within each genotype, with Pearson correlation coefficients of R = 0.71 (P < 1.8 × 10^-6^) for WT replicates and R = 0.78 (P < 4.5 × 10^-9^) for YY1 KO replicates (Fig. S2A). A heatmap of pairwise Pearson correlations with hierarchical clustering confirmed tight grouping of biological replicates within genotypes while clearly separating WT from YY1 KO pro-B samples (Fig. S2B). Consistent with these findings, box plots of log₂-transformed miRNA expression values revealed systematic differences in miRNA expression levels between YY1 WT and YY1 KO pro-B cells, with individual miRNAs shown as overlaid data points (Fig. S2C). Collectively, these analyses establish high technical reproducibility of the miRNA-seq data and demonstrate that YY1 deletion induces robust and consistent alterations in the miRNA expression profile in pro-B cells.

### Comparative miRNA expression profiles in WT and YY1 KO pro-B cells

Our mRNA expression results above indicated that YY1 knockout pro-B cells show reduced expression of some B lineage pathway genes in parallel to increased expression of non-B hematopoietic lineage genes. Given that miRNAs serve as critical post-transcriptional regulators that function by binding to complementary sequences of target mRNAs [43], we set out to investigate whether YY1 regulates miRNA biogenesis and expression in pro-B cells. We identified a total of 35 and 38 known miRNAs expressed in WT or YY1 KO pro-B cells, respectively. The lengths of these miRNAs varied from 18-25 nt (Fig. 3A). There was substantial overlap of 28 miRNA families expressed in both WT and YY1 KO pro-B cells (Figure 3B). Seven miRNAs were detected only in YY1 WT pro-B cells, whereas ten miRNAs were detected only in YY1 KO (Fig. 3B, left and right named miRNAs, respectively). Among the shared miRNAs in WT and YY1 KO pro-B cells, let7d-5p, miR-139-5p, miR-30e-5p, and miR-24-3p showed higher expression in YY1 KO pro-B cells, whereas miR-191-5p, miR-16-5p, let-7f-5p, and let-7a-5p were expressed higher in WT pro-B cells (Fig 3C). However, most expression differences were small (Fig. 3C). Of the seven miRNAs detected only in YY1 WT pro-B cells, miR-142a-5p was the most abundant (Fig. 3D, bottom; see arrow). Among the ten miRNAs detected only in YY1 KO pro-B cells, miR-15b-5p showed the highest expression (Fig. 3D, top; arrow).

**Figure 3.**
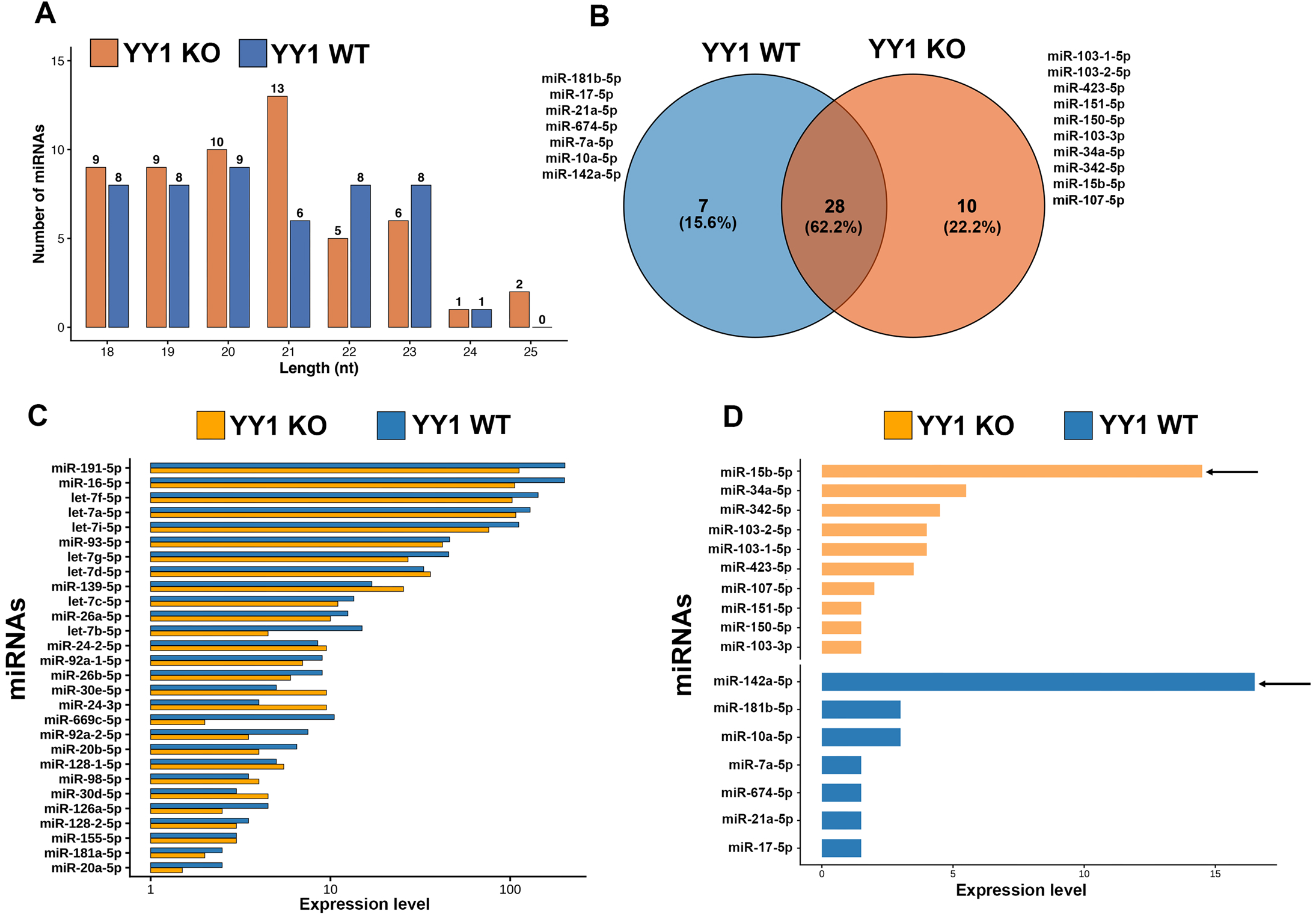
miRNA profiling in YY1 WT and YY1 KO pro-B cells. **(A)** Length distribution of identified conserved miRNAs in YY1 WT (blue bars) and YY1 KO (orange bars) pro-B cells. Bars represent the number of miRNAs identified at each nucleotide length, with the corresponding counts shown above the bars. **(B)** Venn diagram showing the overlap of miRNAs identified in both YY1 WT and YY1 KO pro-B cells, and miRNAs distinctly expressed in either YY1 WT or YY1 KO pro-B cells. Numbers indicate miRNAs unique to each genotype or shared between them. **(C)** Ranked horizontal bars display normalized expression levels of miRNAs commonly detected in both YY1 WT (blue bars) and YY1 KO (orange bars) pro-B cells. **(D)** Expression levels of miRNAs uniquely present in either YY1 WT (blue bars) and YY1 KO (orange bars) pro-B cells. The black arrows highlight the most highly expressed miRNAs in each group (miR-142a-5p in YY1 WT and miR-15b-5p in YY1 KO pro-B cells).

### Target identification and functional analysis of miRNA-regulated transcripts in YY1 WT and YY1 KO pro-B cells

To elucidate the possible impact of the observed miRNA dysregulation after YY1 KO, we performed in silico mRNA target prediction and identified putative mRNA transcripts regulated by the miRNAs that are uniquely expressed in YY1 WT or YY1 KO pro-B cells (Fig. 4A and B). This analysis identified 1,367 and 1,016 predicted target mRNAs expressed in either YY1 WT or YY1 KO pro-B cells, respectively (Fig. 4C, Table S3). Of these, 172 target transcripts (7.8%) were found to be common to both YY1 WT and YY1 KO pro-B cells, while 1195 (54%) and 844 (38.2%) targets were uniquely present in either YY1 WT or YY1 KO samples, respectively (Figure 4C). The miRNAs detected only in YY1 KO pro-B cells showed higher expression in YY1 KO pro-B cells than in YY1 WT pro-B cells (Fig. 4D). To investigate the potential biological functions of these miRNAs expressed in YY1 KO pro-B cells, we used miRDB to identify their predicted mRNA targets and performed KEGG pathway enrichment analysis of these predicted targets. The analysis identified enrichment of pathways involved in B cell development, proliferation, survival, and metabolism, including the PI3K-Akt, mTOR, Wnt, Ras, AMPK, MAPK, cAMP, FoxO, and phosphatidylinositol signaling pathways (Fig. 4E). We then examined the expression of the predicted target genes associated with these pathways. A subset of the predicted targets was significantly downregulated in YY1 KO pro-B cells compared with YY1 WT pro-B cells (Fig. 4F-L). This inverse expression pattern is consistent with potential miRNA-mediated regulation.

**Figure 4.**
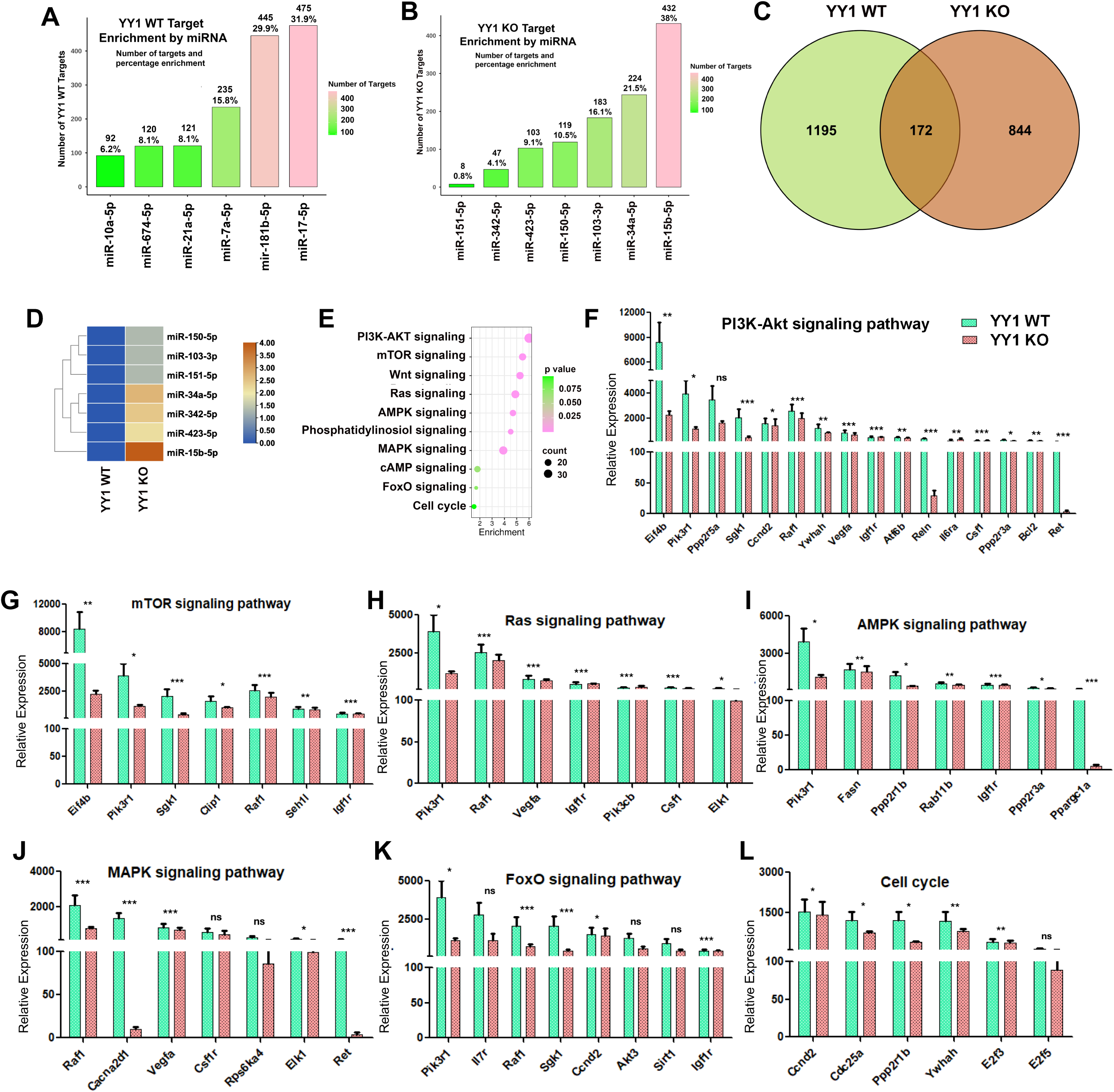
Predicted target transcripts of identified miRNAs and downstream pathway analyses. (A-B) Above each column are shown numbers and percentage enrichment of predicted target genes of miRNAs in WT and YY1 KO pro-B cells. **C)** Venn diagram showing either unique or overlap of miRNA target transcripts in WT and YY1 KO pro-B cells. **(D)** Heatmap representation of normalized miRNA expression levels in YY1 WT and YY1 KO pro-B cells. Blue-to-brown scale shows lower to higher expression, respectively. **(E)** KEGG pathway enrichment analysis of transcripts regulated by miRNAs uniquely present in YY1 KO pro-B cells. **(F-L**) Transcript data show generally reduced expression of individual genes in the various enriched B cell signaling pathways in YY1 KO pro-B cells. All genes associated with each indicated pathway are displayed. Green bars represent YY1 WT transcripts, and pink bars represent YY1 KO transcripts. Asterisks indicate significant differences: P < 0.01, P < 0.001, and P < 0.0001; ns, not significant.

Our target prediction analysis on miRNAs uniquely enriched in YY1 WT pro-B cells showed higher expression in YY1 WT pro-B cells compared to YY1 KO pro-B cells (Fig. S3A) and identified a substantial number of putative mRNA targets (1195 potential targets) (Fig. 4C). KEGG pathway enrichment analysis of these WT-pro-B cell-specific miRNA targets revealed over-representation of multiple signaling pathways crucial for T cells and other alternative lineages, and including Notch, TNF, JAK-STAT, VEGF, and T-cell receptor signaling, as well as genes involved in cytokine-cytokine receptor signaling, and pathways associated with stem cell pluripotency, osteoclast differentiation, and natural killer cell-mediated cytotoxicity (Fig. S3B). These data suggest that elevated expression of these miRNAs in YY1 WT pro-B cells function to repress expression of transcripts typical of alternative lineages, thus serving to maintain B cell identity. However, the statistical significance of the individual RNAs in the pathways controlled by miRNAs specific to YY1 WT pro-B cells was much lower than those in YY1 KO pro-B cells (compare Fig. 4F-L to supplemental Figure S3C-L). Therefore, we chose to focus on the pathways controlled by miRNAs functioning in YY1 KO pro-B cells.

### miR-15b-5p is strongly expressed and is associated with reduced expression of key proliferative and transcriptional networks in YY1 KO pro-B cells

Of the differentially expressed miRNAs we identified here, miR-15b-5p emerged as one of the most strongly and significantly upregulated miRNAs upon YY1 KO (log₂ fold-change = 7.10, adjusted P = 5.48 × 10⁻⁷) (Fig. 5A, Table S4). Target gene enrichment analysis of miR-15b-5p revealed strong over-representation of pathways involved in cell proliferation, cell cycle progression, regulation of apoptosis, and transcriptional control (Fig. 5B). These processes are critically important during the highly proliferative pro-B cell stage, where tight regulation of survival and division is essential for proper B cell development. Consistent with these findings, transcription factor enrichment analysis identified several master regulators of these processes as major targets of miRNA15b-5p. Most notably BCL6B and multiple members of the E2F transcription factor family (E2F1, E2F2, E2F3, E2F4, and E2F7) were identified (Fig. 5C). BCL6B is a critical transcriptional regulator in lymphoid cells, while the E2F transcription factors are central drivers of cell cycle progression and proliferation processes that are highly active during the pro-B to pre-B cell transition.

**Figure 5.**
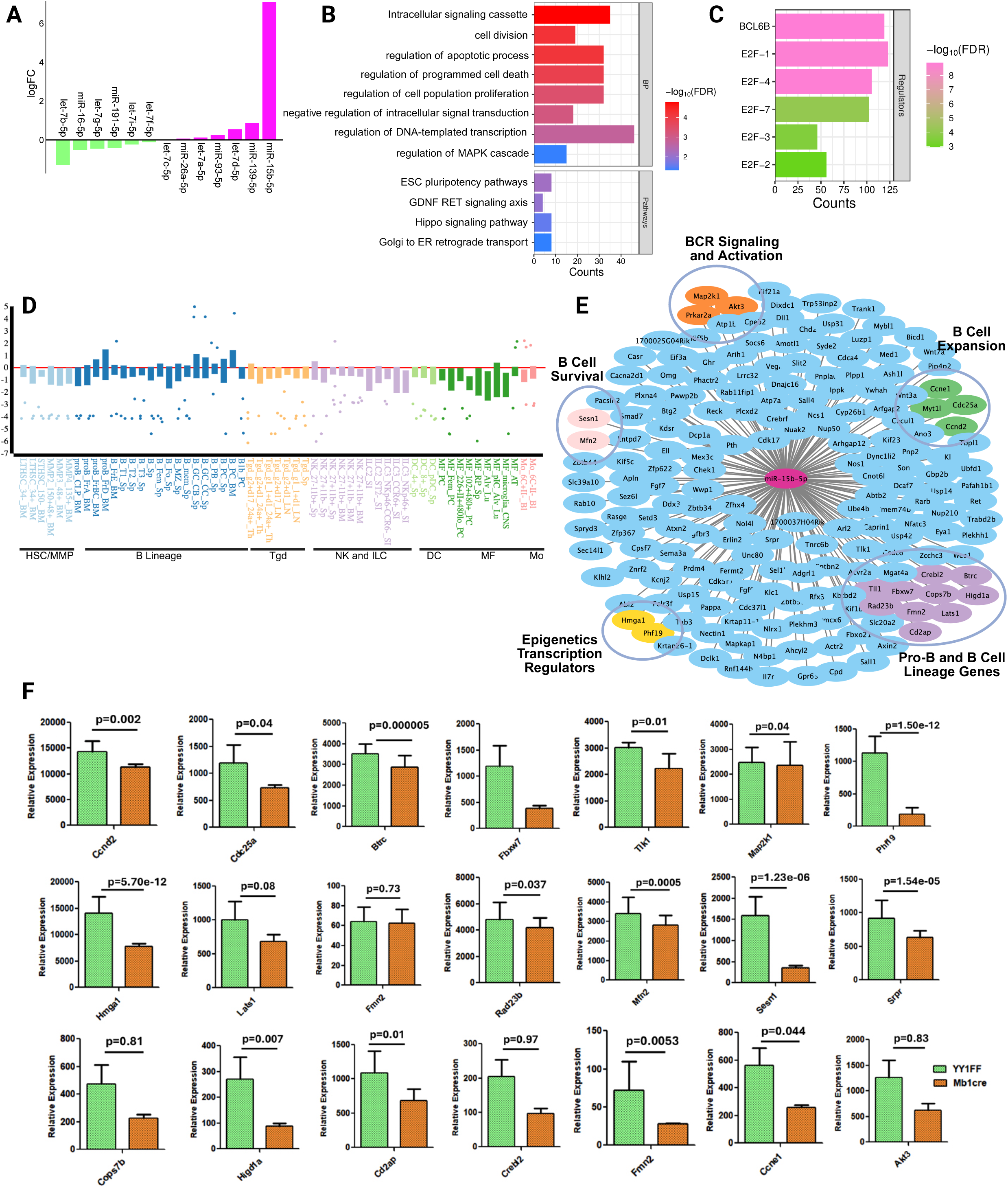
miR-15b-5p targets key proliferative and transcriptional networks in pro-B cells. **(A)** Bar graph plot showing differential miRNA expression in YY1 WT (green bars) versus KO (pink bars) pro-B cells. miR-15b-5p is among the most strongly and significantly expressed miRNAs in pro-B cells. **(B)** biological process enrichment and **(C)** transcription factor target enrichment of predicted miR-15b-5p targets **(D)** ImmGen expression atlas of miR-15b-5p target mRNAs demonstrate expression levels at the pro-B cell stage, with additional peaks at the germinal center and plasma cell stages which represent other B lineage stages with high proliferative capacity. **(E)** miR-15b-5p target network analysis showing the schematic summary of major functional B lineage categories enriched among miR-15b-5p target mRNAs. These processes encompass BCR signaling and activation, B cell expansion, pro-B and B cell lineage genes, epigenetic and transcription regulators, and B cell survival genes. **(F)** Expression profiles of key miR-15b-5p target mRNAs. Data are shown as mean ± standard deviation from three independent biological replicates. Statistical p values are shown above each graph

The ImmGen expression atlas showed that miR-15b-5p target mRNAs exhibit peak expression within the pro-B and early B cell compartments and are poorly expressed in other hematopoietic cell types (Fig. 5D). The miR-15b-5p target RNAs are also strongly expressed across multiple stages of early B lymphopoiesis, including stages involving BCR signaling and activation, B cell expression of pro-B cell lineage genes, activation of epigenetic regulators, and control of B cell survival (Fig. 5E). RNA transcript profiling revealed significant downregulation of multiple miR-15b-5p target genes in YY1 KO pro-B cells compared with YY1 WT pro-B cells (Fig. 5F). Collectively, these data indicate that elevated miR-15b-5p expression in YY1 KO pro-B cells reduces key proliferative and anti-apoptotic networks orchestrated by BCL6B and E2F transcription factors and disrupts multiple B cell pathways and functions needed for developmental progression of B-lineage cells, thus providing opportunity for potential alternative lineage development.

### YY1 KO disrupts B-cell commitment and promotes lineage plasticity coincident with increased miRNA-15b-5p expression

We previously showed that YY1 KO pro-B cells, but not YY1 WT pro-B cells, grown on OP9-DL4 feeder cells for two weeks, were capable of yielding scRNA-seq profiles indicative of alternative hematopoietic lineages, including cells with stem cell transcript profiles [2]. High quality scRNA-seq data was generated from WT and YY1 KO pro-B cells directly isolated from mice, or after incubation of these cells for two weeks on OP9-DL4 feeders (Fig. S5A and B). The scRNA-seq UMAPs from WT and YY1 KO pro-B cells directly isolated from mice showed very similar UMAP profiles (Fig. S5A and B, panels 3 and 4). However, when these WT and YY1 KO pro-B cells were incubated on OP9-DL4 feeders for two weeks, very distinct results were obtained. While YY1 WT pro-B cells maintained their B lineage phenotype (Fig. S5A and B, panel 2), YY1 KO pro-B cells nearly completely lost their B lineage phenotype and adopted RNA expression patterns indicative of dendritic cells, monocytes, macrophages, and even stem cells (Fig. S5A and B, panel 1).

We hypothesized that the predicted mRNA targets of miRNA-15b-5p including Hmga1, Lats1, Mfn2, Mybl1, Sesn1, Btrc, Fbxw7, Akt3, and Ops7b, would show reduced expression in YY1 KO pro-B cells after two weeks of culture on OP9-DL4 feeder cells. To examine this possibility, we evaluated scRNA-seq data from pro-B cells isolated directly from YY1 WT or YY1 KO pro-B mice, or after growth for two weeks on OP9-DL4 cells. Interestingly, we found that expression of Hmga1, Lats1, Mfn2, Mybl1, Sesn1, Btrc, Fbxw7, Akt3, and Cops7b mRNAs were indeed very dramatically reduced in YY1 KO pro-B cells grown for 2 weeks on OP9-DL4 feeder cells compared to all other conditions (Fig. 6A, compare panel 1 to panels 2-4, and Fig. 6B). This inverse expression pattern is consistent with miR-15b-5p-mediated regulation.

**Figure 6.**
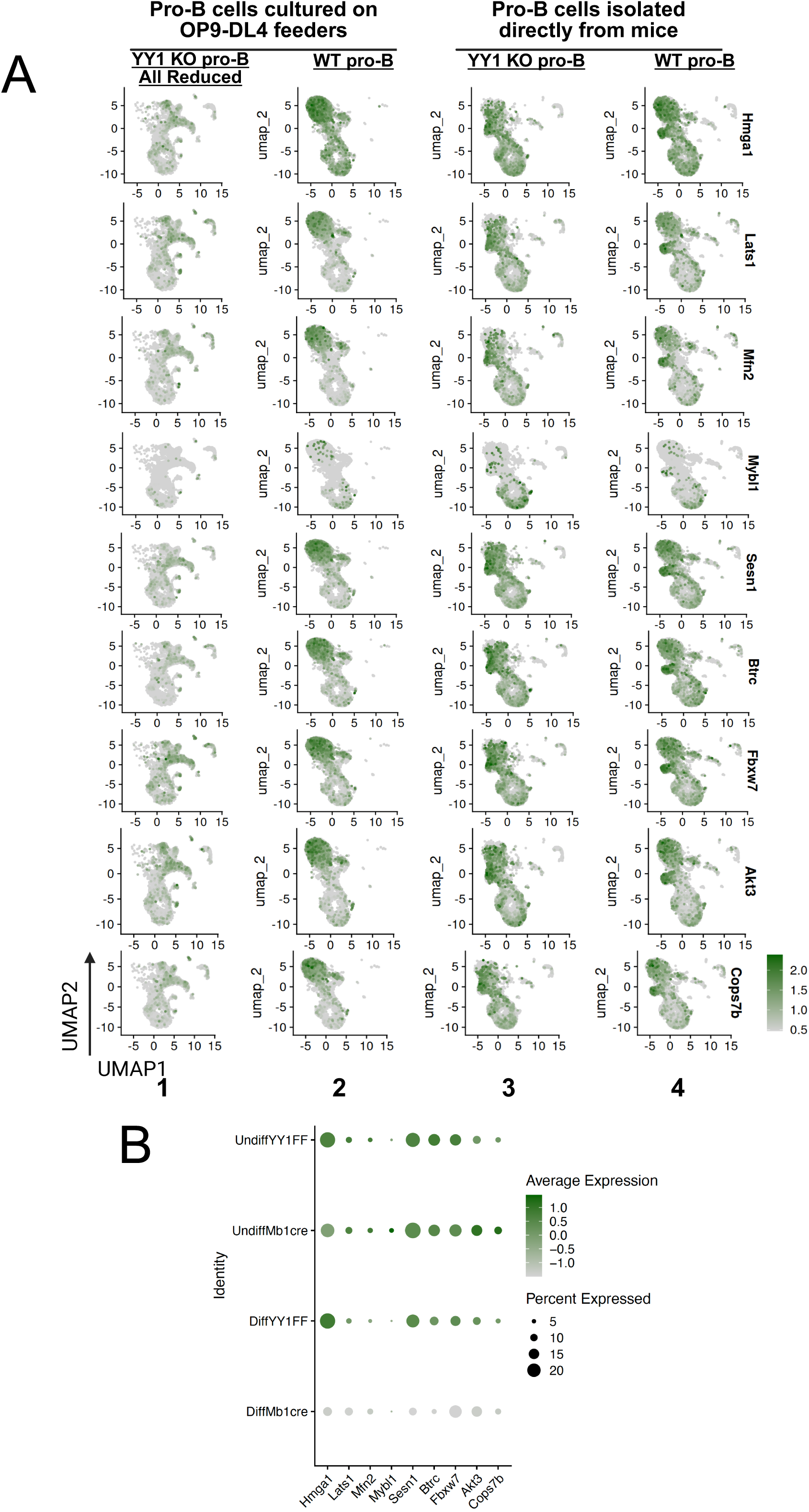
mRNAs targeted by miR-15b-5p are down-regulated in YY1 KO pro-B cells grown on OP9-DL4 feeders. **(A)** WT and YY1 KO pro-B cells were isolated directly from mice and either subjected to scRNA-seq evaluation (vertical panels 3 and 4), or grown on OP9-DL4 feeders in the presence of IL7, SCF, and Flt3L for 14 days prior to scRNA-seq evaluation (vertical panels 1 and 2). miR-15b-5p target transcripts in YY1 KO pro-B cells were dramatically reduced after growth on OP9-DL4 feeders (vertical panel 1). **(B)** Quantitation of target mRNA expression. Dot color represents the scaled average expression, and dot size represents the percentage of cells expressing each gene within the indicated condition on the y axis. Transcript identifies are shown on the x axis.

Finally, we performed pseudotime trajectory analysis of scRNA-seq data from YY1 KO pro-B cells cultured on OP9-DL4 feeders for 14 days to infer the transcriptional relationships of cells classified as stem/progenitor-like cells progressing toward alternative hematopoietic lineages (Fig. 7A and B). Cells classified by SingleR as stem/progenitor-like were located in the lower region of the UMAP plot, whereas cells classified as alternative hematopoietic cell types were located primarily in the upper region (Fig. 7A; see arrows). Pseudotime analysis inferred a continuous and highly branched organization of transcriptional states rather than a simple binary bifurcation (Fig. 7B). Using the progenitor/stem-like compartment as the pseudotime root, the inferred trajectory extended through a central region and separated into branches associated with monocyte, macrophage, dendritic-cell, and mast-cell transcriptional states. A relatively small subset of cells retained a B-lineage transcriptional identity and was positioned apart from the predominant inferred trajectories, suggesting limited maintenance of B lineage identity under these culture conditions. Pseudotime gradients were associated with alternative hematopoietic transcriptional states, suggesting an organized pattern of cell-state variation. The presence of several branching points indicates that YY1 KO cells occupy a broad range of transcriptional states associated with multiple non-B-cell identities rather than a single alternative state. Together, these findings support a model in which YY1 is critical for the maintenance B cell identity. In the absence of YY1, but under Notch signaling conditions, pro-B cells exhibit reduced lineage fidelity and increased transcriptional plasticity associated with alternative hematopoietic cell states.

**Figure 7.**
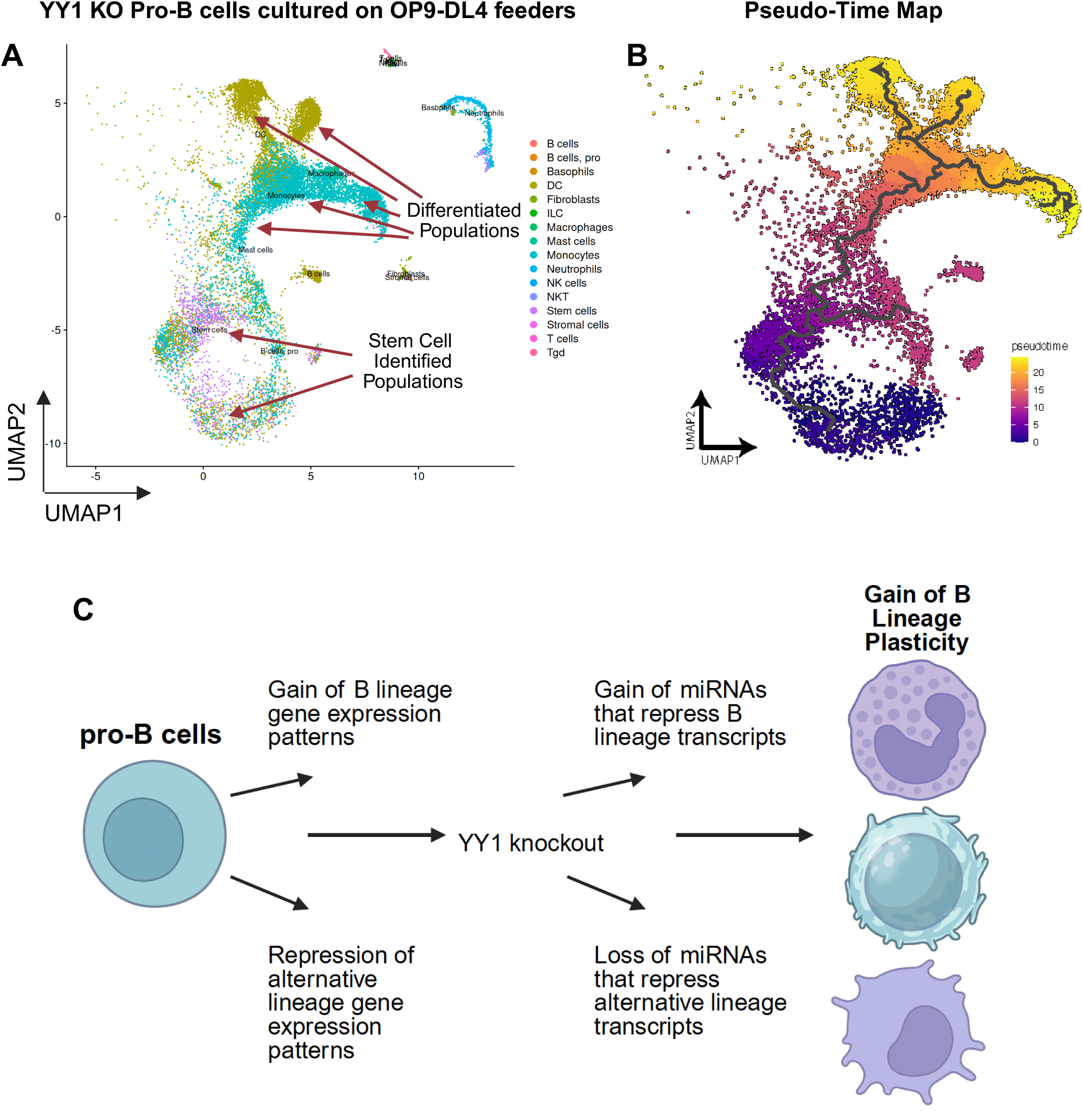
YY1 KO disrupts B cell lineage commitment and drives cell fate toward alternative hematopoietic lineages. **(A)** UMAP projection of integrated single-cell RNA-seq data from YY1 KO pro-B cells cultured on OP9-DL4 feeders with IL7, SCF, and Flt3L for 2 weeks. Arrows indicate SingleR defined cell type identification for stem cells, dendritic cells, macrophages, monocytes, as well as other cells. **(B).** Pseudo-Time UMAP pathways starting at stem cells and progressing toward various hematopoietic cell types. YY1 KO pro-B cells were grown 2 weeks on OP9-DL4 feeders in IL7, SCF, and Flt3L. RNA was isolated and subjected to scRNA-seq analyses. The black lines trace differentiation from stem cells into cells acquiring the transcript phenotypes of various hematopoietic lineages. **(C)** Model of the impact of YY1 KO on gain of miRNAs that reduce B lineage transcripts, and the loss of miRNAs that result in increased alternative lineage transcripts.

## Discussion

YY1 has long been recognized as a multifunctional transcription factor involved in chromatin organization, enhancer-promoter communication, and lineage-specific gene regulation [8, 44]. Yet, its precise role in maintaining early B cell identity has remained incompletely understood. In this study, using integrated mRNA-seq, miRNA-seq, and single-cell transcriptomic analyses, we explored the role of YY1 as a central regulator of transcriptional fidelity and lineage commitment in pro-B cells. Our data provide evidence that YY1 simultaneously promotes B-lineage gene expression programs while repressing alternative hematopoietic fates through coordinated transcriptional and post-transcriptional mechanisms.

Global transcriptomic profiling revealed extensive reprogramming upon YY1 loss, with more than 1,233 genes differentially expressed and a clear separation between WT YY1 and YY1 KO pro-B cells. Notably, genes downregulated in YY1 KO pro-B cells were strongly enriched for canonical B cell pathways, including B cell receptor signaling and lymphocyte activation, consistent with prior reports demonstrating the requirement of YY1 in immunoglobulin locus contraction and B cell development [2, 16, 45, 46]. In contrast, the upregulated gene programs were enriched for cytokine signaling, JAK-STAT, NF-κB, and hematopoietic lineage pathways, indicating activation of transcriptional programs normally repressed in pro-B cells. This dual regulatory role supports the concept that YY1 functions not only as a transcriptional activator of lineage-specific genes but also as a gatekeeper that suppresses alternative lineage potential, consistent with its known role in transcriptional repression and epigenetic regulation [47, 48].

While we previously showed YY1 KO is linked to transcriptional and developmental plasticity [2], our findings here reveal that miRNA-mediated regulation can also play a role in this loss of lineage identity. Rather than simply blocking B cell differentiation, YY1 loss creates a permissive transcriptional state that enables activation of alternative lineage programs, including myeloid and inflammatory gene modules. In this context, altered miRNA networks provide an additional layer of regulation that may further destabilize B cell identity upon YY1 KO and facilitate activation of alternative lineage programs (Fig. 7C). This phenotype is particularly evident in single-cell RNA-seq analyses of miRNA targets, where YY1-deficient pro-B cells exhibit marked heterogeneity and bifurcation into non-B lineages under Notch-driven conditions. These results extend previous observations of lineage plasticity in hematopoietic progenitors and suggest that YY1 is essential for stabilizing lineage trajectories during early B lymphopoiesis as well as in other lineages [2, 6, 7].

Beyond transcriptional control, our findings uncover a substantial role for YY1 in regulating miRNA expression landscapes. miRNA profiling identified both shared and genotype-specific miRNAs, with distinct regulatory networks associated with YY1 presence or absence. miRNAs enriched in YY1 KO pro-B cells preferentially targeted signaling pathways critical for B cell development and survival, including PI3K-Akt, MAPK, Wnt, and mTOR pathways. Downregulation of these target genes after YY1 KO is consistent with established models of miRNA-mediated repression, in which elevated miRNA levels destabilize transcripts and suppress translation [21, 49]. Conversely, miRNAs enriched in WT pro-B were predicted to target genes associated with T-cell and alternative lineage differentiation pathways, suggesting that YY1 maintains lineage commitment in part by establishing a miRNA environment that suppresses inappropriate transcriptional programs.

Among miRNAs studied here, miR-15b-5p emerged as a critical regulator downstream of YY1. This miRNA was one of the most strongly upregulated candidates in a YY1 KO pro-B background and is predicted to target key regulators of proliferation and transcription, including BCL6B and multiple E2F family members. E2F transcription factors are well-established regulators of cell cycle progression and are essential for the proliferative expansion of early B cell progenitors [50–53], while BCL6 family proteins play important roles in lymphocyte differentiation and survival [54]. The coordinated downregulation of these targets in YY1 KO pro-B cells suggests that miR-15b-5p contributes to impaired proliferative capacity and disrupted developmental progression. Furthermore, the enrichment of miR-15b-5p target genes in pro-B and early B cell compartments, as shown by ImmGen data, supports a stage-specific regulatory role. These findings are consistent with previous studies implicating miR-15 family members in cell cycle regulation and hematopoietic differentiation [55, 56].

The single-cell RNA-seq expression pattern of miR-15b-5p target transcripts, including Hmga1, Lats1, Mfn2, Myb1l, Sesn1, Btrc, Akt3, Cops7b, and Fbxw7, showed markedly reduced expression profiles in YY1 KO pro-B cells cultured on OP9-DL4 feeders (Fig. 6), consistent with enhanced miRNA-mediated repression. Further, pseudotime analysis demonstrated that YY1 KO pro-B cells grown on OP9-DL4 feeders do not follow a B cell lineage trajectory but instead traverse a branched landscape with multiple developmental endpoints, including monocyte/macrophage, and DC cell lineages. This widespread trajectory diversification indicates that YY1 is essential for suppressing multilineage potential, and to maintain directional progression through the B cell lineage.

In conclusion, we found YY1 is a central regulator of B cell lineage commitment that integrates transcriptional and post-transcriptional regulatory mechanisms (Fig. 7C). While previous studies have established a role for YY1 in maintaining chromatin architecture and lineage stability [2], our findings extend this model by revealing that YY1-dependent lineage stabilization involves both transcriptional regulation and miRNA-mediated post-transcriptional control. Together, these findings support a model in which YY1 establishes a chromatin and transcriptional environment that preserves B cell identity, while YY1-regulated miRNA networks reinforce this lineage-restricted state by controlling expression of key developmental regulators (Fig. 7C). In particular, miR-15b-5p emerges as a key downstream effector of YY1 loss, targeting multiple B-lineage genes and reinforcing their repression, linking YY1 loss to transcriptional plasticity. Our study reveals how miR-15b-5p driven regulation contributes to the destabilization of B cell identity and the emergence of lineage plasticity. These findings uncover a new layer of regulation through which miRNAs contribute to lineage instability and highlight their importance in shaping cell fate decisions during hematopoietic development.

**Table 1.** Summary of RNA-seq read mapping statistics for wild-type and YY1-deficient pro-B cells. Mapping statistics of high-throughput RNA sequencing data from wild-type (YY1FF) and YY1-knockout (Mb1cre) pro-B cells across biological replicates. Raw reads represent the total number of sequencing reads obtained. After filtration, the remaining reads are subjected to quality control (Phred score > 30) and adaptor trimming. Mapped and unmapped reads show the number and percentage of reads successfully aligned to the reference genome (mm10) or remaining unaligned, respectively.

| **Samples** | **Raw reads** | **After filtration (q**<br>**Value >30 &**<br>**adaptor)** | **Mapped** | **Unmapped** |
| --- | --- | --- | --- | --- |
| YY1FF1 | 13719318 | 13719318 | 1624605<br>(11.842%) | 12094713<br>(88.158%) |
| YY1FF2 | 12268534 | 12226491 | 1023959<br>(8.375%) | 11202532<br>(91.625%) |
| Mb1cre1 | 12799905 | 12799905 | 1487815<br>(11.624%) | 11312090<br>(88.376%) |
| Mb1cre2 | 13783625 | 13783625 | 1146116<br>(8.315%) | 12637509<br>(91.685%) |

