## Supplemental Figures 1 to 5 for "miRNA-mediated regulation of lineage plasticity in YY1 knockout pro-B cells"

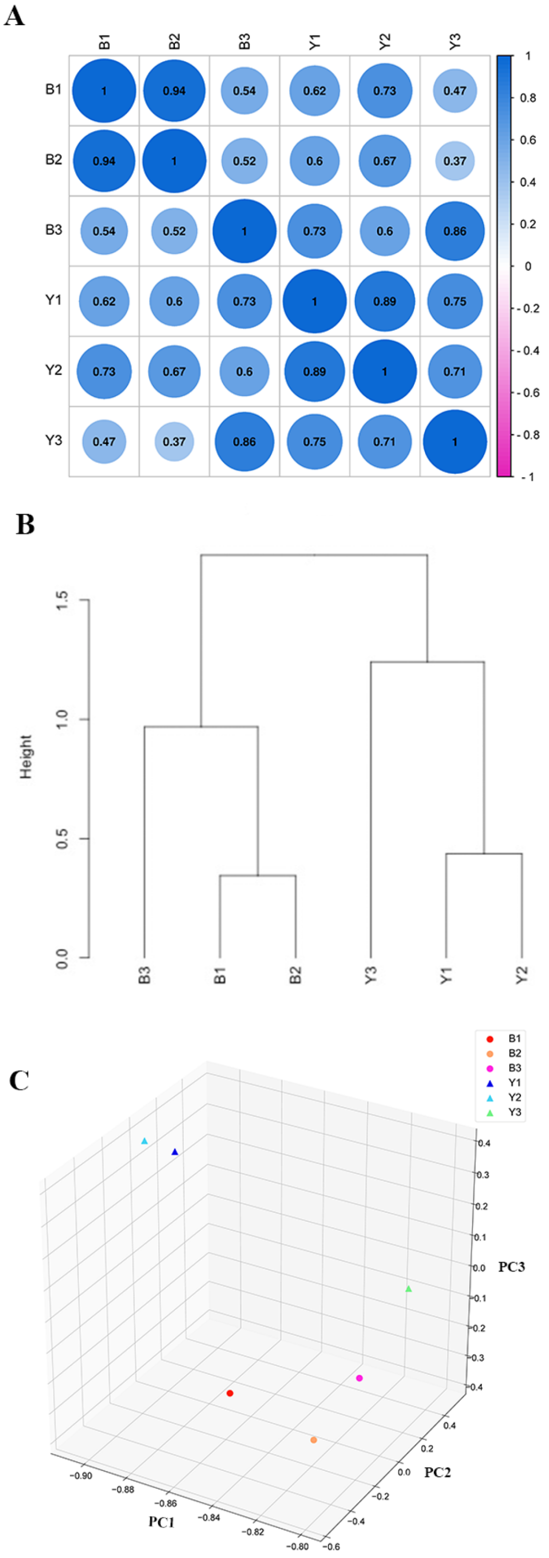


Bano et al Figure S1

**Supplementary Figure S1. Quality control and reproducibility assessment of RNA-seq data from WT and YY1 KO pro-B cells.**

**(A)** Pearson correlation matrix showing pairwise correlation coefficients among biological replicates. Samples B1-B3 represent YY1 WT pro-B cells, and Y1-Y3 represent YY1 KO pro-B cells. Color intensity and circle size indicate the strength of the correlation (r=1 indicating perfect positive correlation). B1 and B2 were strongly correlated, whereas B3 showed lower correlations with the other WT replicates.

**(B)** Hierarchical cluster dendrogram based on global gene expression profiles, demonstrating the grouping of biological replicates. These analyses show despite the lower pairwise correlations of B3 with B1 and B2, the WT and YY1 KO samples formed distinct major branches.

**(C)** Principal component analysis illustrating the clustering and separation of YY1 WT and YY1 KO biological replicates.

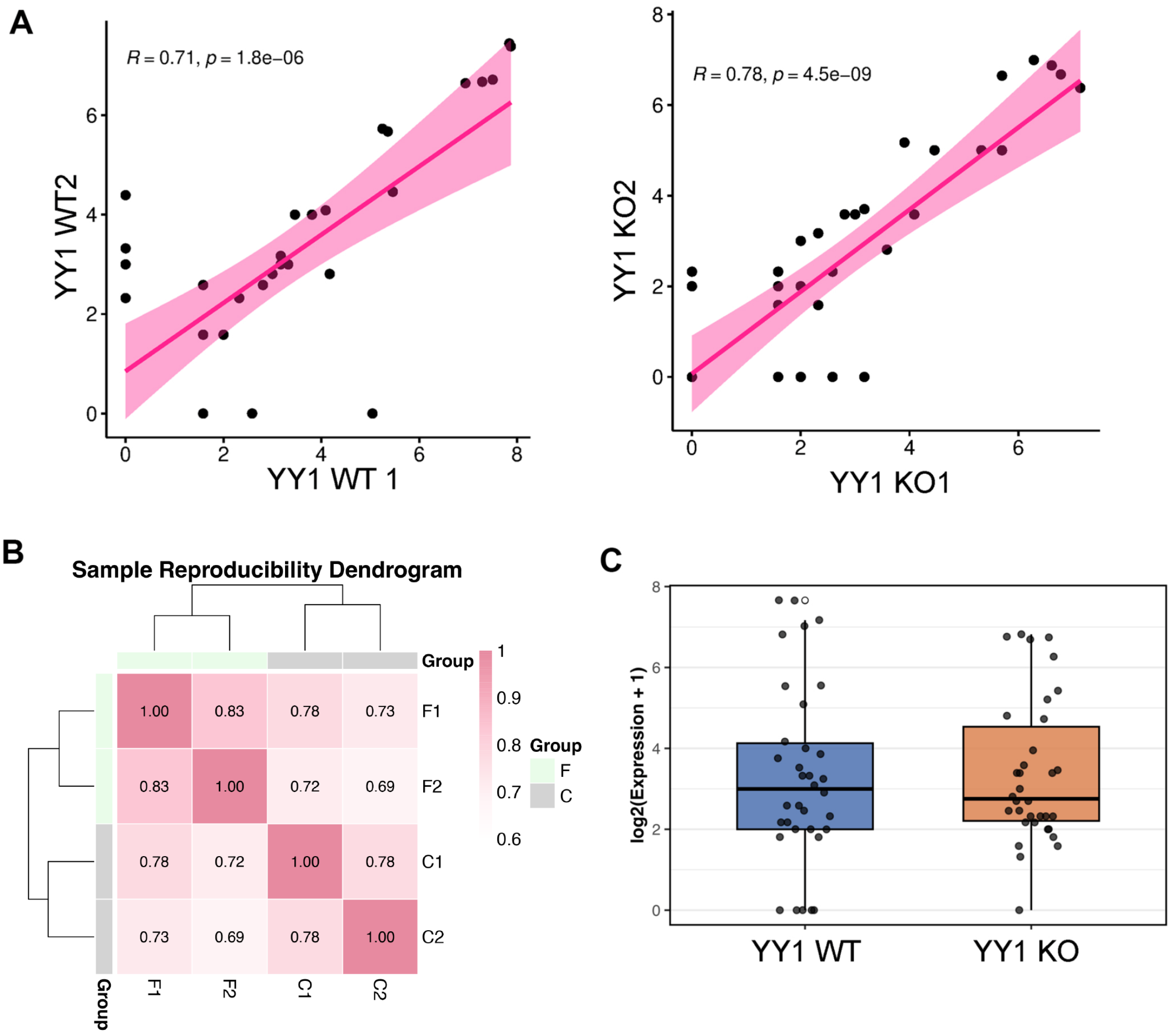


Bano et al

Figure S2

**Supplementary Figure S2. Reproducibility assessment of miRNA-seq biological replicates.**

**(A)** Scatter plots of $log_2$-transformed normalized miRNA expression values showing strong correlations between biological replicates of the same genotype. Left panel: wild-type pro-B cells (YY1WT1 versus YY1WT2). Right panel: YY1 KO pro-B cells (YY1 KO1 versus YY1 KO2). Pearson's correlation coefficient (r) and the associated two-tailed P-value are displayed for each comparison.

**(B)** Heatmap displaying pairwise Pearson correlation coefficients across all samples. Samples are labeled as follows: F1 and F2 (YY1WT1 and YY1WT2) and C1 and C2 (YY1KO1 and YY1KO2). The accompanying dendrogram (hierarchical clustering) illustrates the grouping of biological replicates by genotype. Color intensity represents the strength of the correlation, with dark pink indicating high positive correlation and light pink indicating a lower correlation. These analyses confirm high reproducibility within genotypes and clear separation of miRNA expression profiles between wild-type and YY1 KO pro-B cells.

**(C)** Box plots showing the distribution of $log_2$-transformed miRNA expression values in YY1 wild-type (YY1 WT) and YY1 knockout (YY1 KO) pro-B cells. The plot highlights the overall shift in miRNA expression upon YY1 deletion, consistent with the clear genotype separation observed in the correlation analyses.

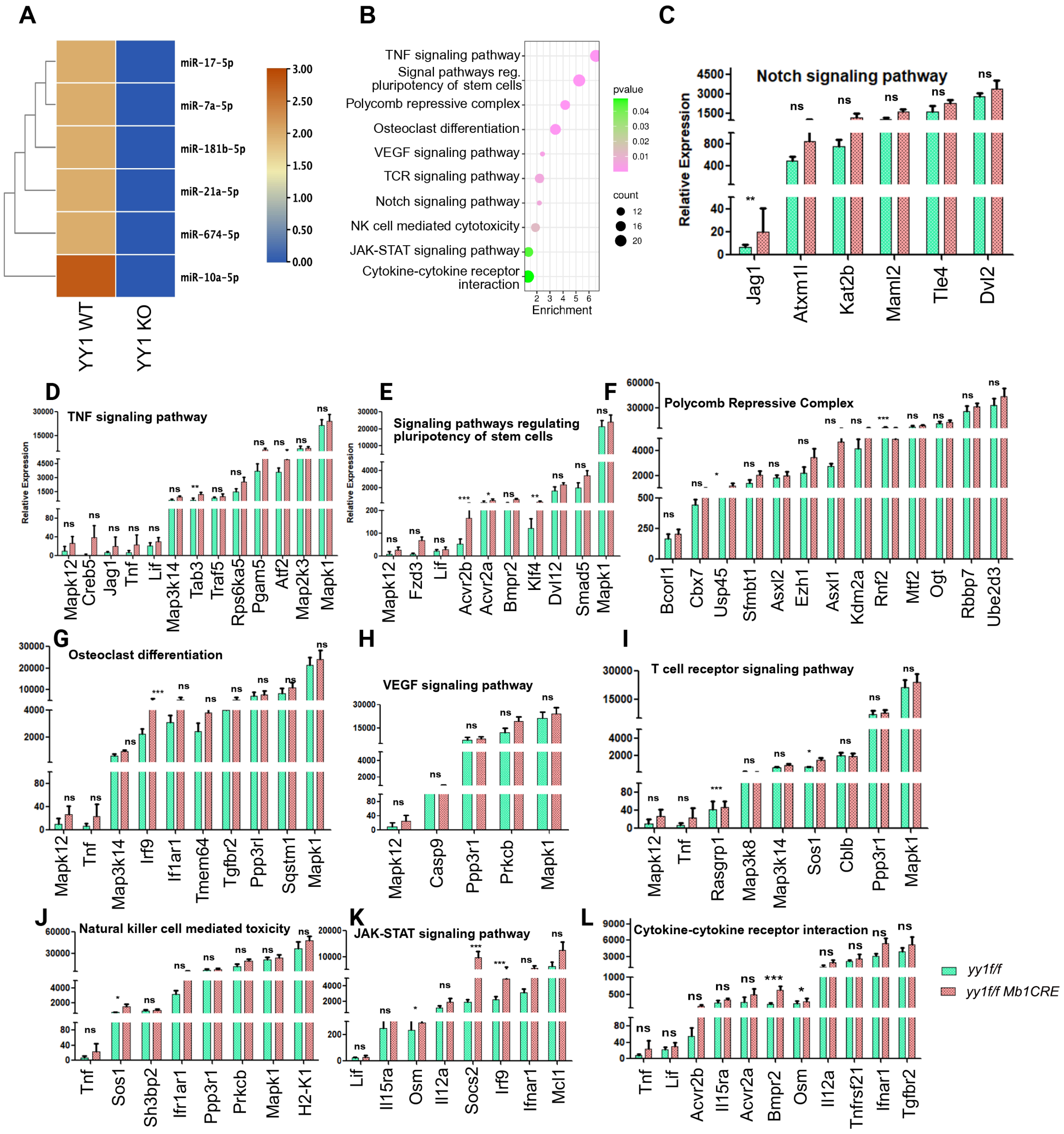


Bano et al    Figure S3

**Figure S3. KEGG pathway enrichment of YY1 WT-specific miRNA targets and expression profiling of associated transcripts.**

**(A)** Heatmap showing normalized expression levels of miRNAs in YY1 WT and YY1 KO pro-B cells. The blue-to-brown color scale represents low to high expression, respectively.

**(B)** KEGG pathway enrichment analysis of target transcripts regulated by miRNAs uniquely expressed in YY1 WT pro-B cells.

**(C-L)** mRNA expression levels of genes involved in key signaling and cellular pathways that are targeted by miRNAs expressed in WT pro-B cells. **(C)** Notch signaling, **(D)** TNF signaling, **(E)** Signaling pathways regulated to stem cell pluripotency, **(F)** Polycomb repressive complex, **(G)** Osteoclast differentiation, **(H)** VEGF signaling, **(I)** T cell receptor signaling, **(J)** Natural killer cell-mediated toxicity, **(K)** JAK-STAT signaling, and **(L)** Cytokine–cytokine receptor interaction. Data are shown as mean ± standard deviation from three independent biological replicates. Asterisks indicate statistically significant differences (*P < 0.01, **P < 0.001, ***P < 0.0001).

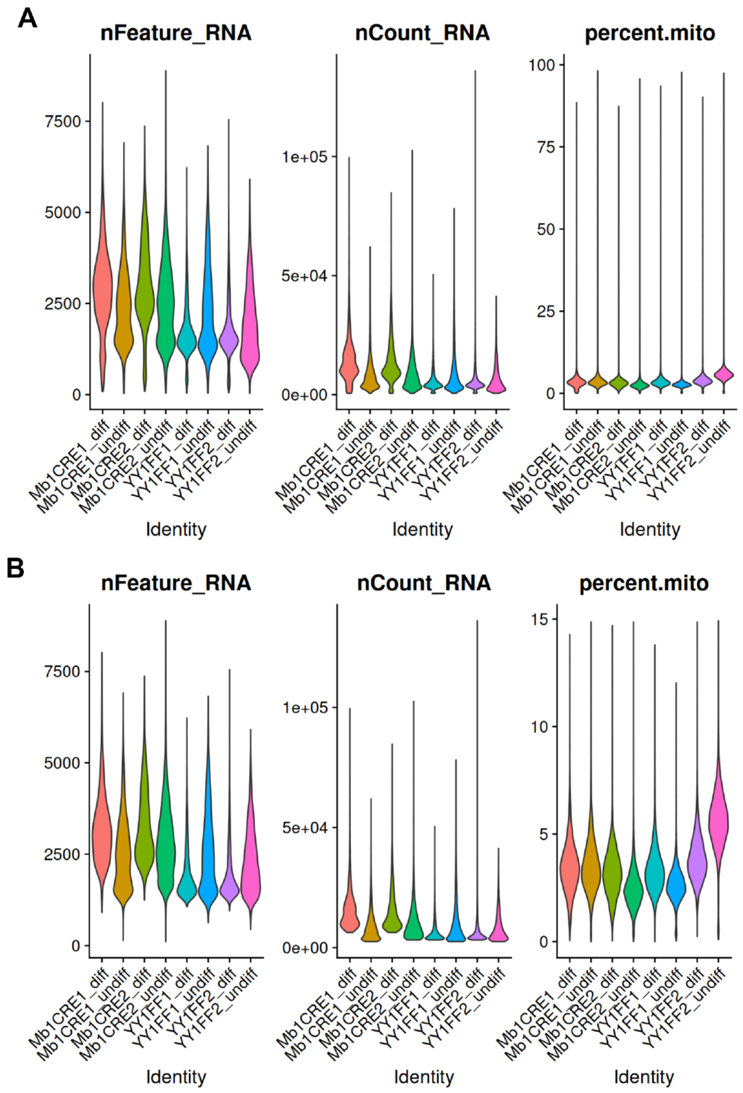


Bano et al Figure S4

**Figure S4. Quality control and filtering of single-cell RNA-seq data.**

**(A)** Violin plots showing the distribution of key quality metrics across cell identities before filtering: number of detected genes (nFeature_RNA), total RNA counts (nCount_RNA), and percentage of mitochondrial transcripts (percent.mito).

**(B)** The same quality metrics after stringent filtering, demonstrating effective removal of low-quality cells while maintaining the integrity of major cell populations.

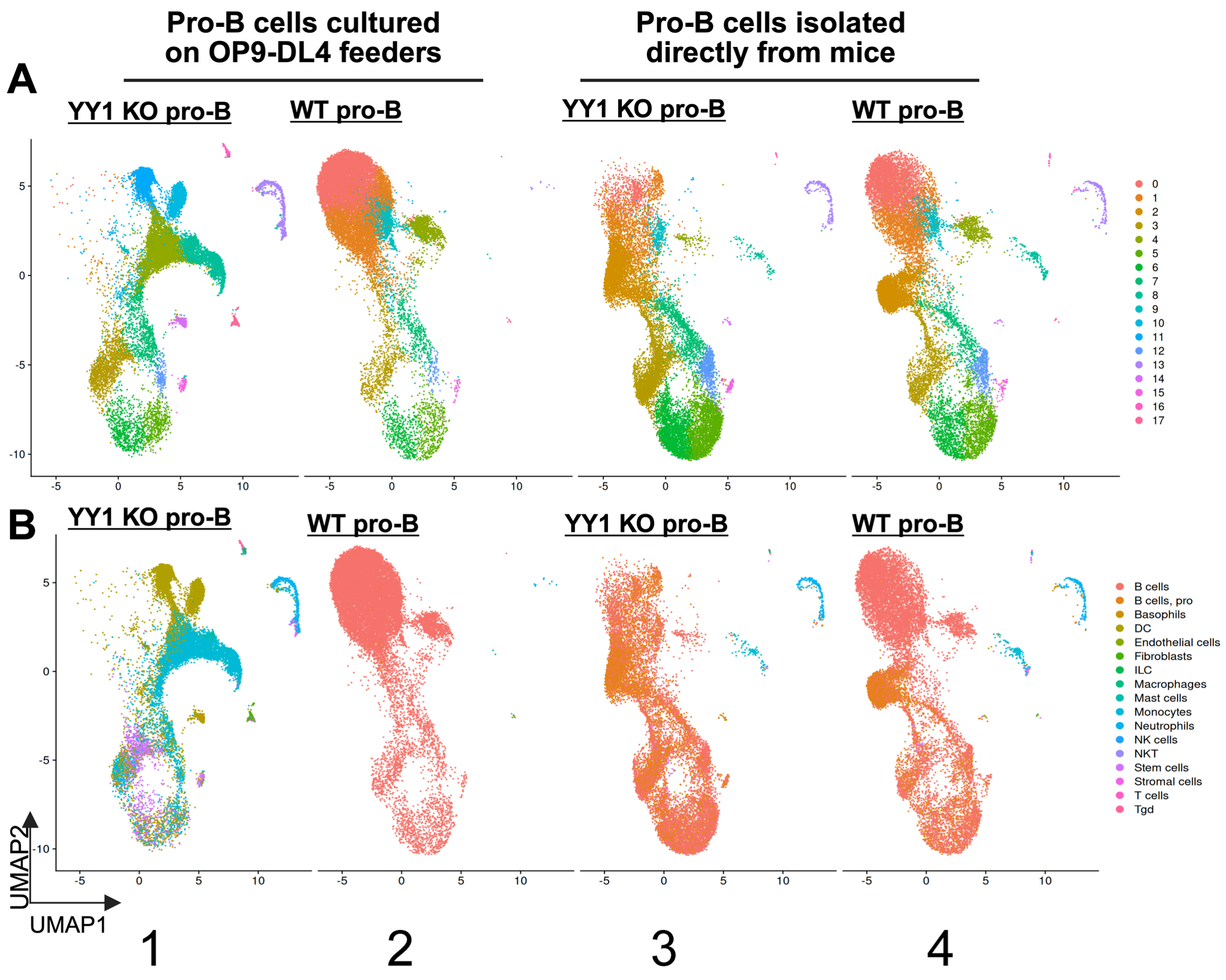


Bano et al Figure S5

**Figure S5. YY1 KO pro-B cells are poised to adopt distinct hematopoietic cell types.**

YY1 WT and YY1 KO pro-B cells were grown on OP9-DL4 feeders in the presence of IL7, SCF, and Flt3L for 14 days. Cells were then subjected to single cell RNA sequence analyses.

**(A)** UMAP profiles of scRNA-seq data from YY1 KO pro-B cells or WT pro-B cells grown on OP9-DL4 feeders for two weeks (Panels 1 and 2) or the same cells directly isolated from mice (panels 3 and 4).

**(B)** Single R identification of cell types deduced from the scRNA-seq data in (A) indicate that YY1 KO pro-B cells on OP9-DL4 feeders for two weeks adopt the phenotypes of numerous alternative hematopoietic lineage cell types. The right panel provides the color key for individual cell types.
